# A Simple Generative Model for the Prediction of T-Cell Receptor - Peptide Binding in T-cell Therapy for Cancer

**DOI:** 10.1101/2025.03.18.643937

**Authors:** Athanasios Papanikolaou, Vladimir Sivtsov, Enrica Zereik, Eliana Ruggiero, Chiara Bonini, Fabio Bonsignorio

## Abstract

**Objective:** To develop a deep learning model capable of predicting epitope peptides recognized by specific CDR3 (Complementarity-Determining Region 3) sequences of T-cell receptors (TCRs) in the context of Major Histocompatibility Complex (MHC) molecules, addressing the challenges of incomplete datasets and the need for novel sequence generation in adoptive T-cell therapy for cancer.

**Methods:** We implemented a sequence to sequence generative model named “GRIP” (Generative Reconstruction of antIgen Peptides) using a Long Short-Term Memory (LSTM) network with attention mechanisms. The model was trained and validated on publicly available datasets, employing data balancing, label smoothing, and dynamic learning rate scheduling to enhance performance and generalization. Accuracy was assessed at the amino acid level.

**Results:** The model achieved a training accuracy of 97% and a test accuracy of 85% for predicting epitope sequences at the amino acid level. Probabilistic sequence generation allowed GRIP to produce biologically plausible epitope sequences, even for unseen CDR3 inputs. Attention-based interpretability provided insights into the model’s focus on critical sequence elements. The model outperformed existing approaches in handling data imbalance and generalization to novel epitopes.

**Conclusion:** GRIP offers a novel solution to the TCR-epitope binding problem by generating potential epitope sequences instead of matching to known data, addressing a fundamental gap in existing models. This approach has significant implications for personalized immunotherapy, facilitating the design of targeted T-cell therapies for cancer.

## 1. Introduction

The immune system’s ability to target and eliminate pathogens and cancer cells is a cornerstone of effective immunotherapy. Central to this process is the precise interaction between epitope peptides, CDR3 sequences of TCRs, and MHC molecules. As discussed in [1], these interactions are fundamental in determining the immune system’s recognition and response to antigens, which is crucial in activating T-cells. We will explore these terms in more detail later in the dataset subsection.

However, predicting which epitope peptides can effectively bind to specific CDR3 sequences and MHC molecules remains a significant challenge. Traditional experimental methods for identifying these interactions are labour-intensive, costly, and time-consuming [2]. Moreover, the vast diversity of potential epitopes and the specificity of TCR-MHC interactions further complicate this task.

There is a growing need for computational models to address these challenges that can accurately and efficiently predict the interactions between epitope peptides, CDR3 sequences, and MHC molecules. Such models would significantly advance personalized immunotherapy, allowing for the design of targeted treatments tailored to individual patients’ immune profiles. This would enhance the efficacy of immunotherapy and reduce the time and cost associated with developing these treatments.

In this study, we present a novel computational approach to predict epitope peptides based on given CDR3 sequences and MHC molecules using deep learning techniques. Our sequence-to-sequence model “GRIP – Generative Reconstruction of antIgen Peptides” leverages recurrent neural networks (RNNs) with LSTM layers, which are particularly effective in capturing the dependencies and patterns within sequential data, and generates “new” epitopes even with unseen CDR3 sequences. As highlighted in [3], RNNs and LSTMs are especially suited for tasks involving temporal or sequential data due to their ability to maintain memory over longer sequences, making them highly applicable for biological sequence analysis. The model also incorporates convolutional layers, which are adept at capturing local patterns within data, and attention mechanisms, which allow the model to focus on the most relevant parts of the input sequences, further enhancing its predictive capabilities.

**Table 1:** Summary.

|  |  |
| --- | --- |
| **Problem** | Predicting which epitope peptides bind to specific CDR3 sequences of T-cell receptors for cancer immunotherapy. |
| **What is Already Known** | Most existing models match CDR3 sequences with known epitope peptides, but struggle with data imbalance and generalizing to unseen epitopes. This limits their applicability in personalized treatments. |
| **What This Paper Adds** | This study introduces a generative LSTM model named GRIP that can predict new epitope sequences for unseen CDR3 inputs, enhancing generalizability and addressing data imbalance through probabilistic sequence generation. |

## 2. Related Work

Many different approaches, mostly relying on deep learning methodologies, have been proposed in the literature. Key points, common to all of the proposed systems, are the attention to system *interpretability* and the problem of biased and insufficient datasets. These two can lead to actual non-explainability of the model behaviour and hide algorithmic errors [4].

Work in [5] discusses the importance of determining the main factors of TCR-epitope interactions at a molecular level, underlining how the 3D structure of the molecule could offer new insights on the binding occurency. To this aim, various methods are trying to model such 3D structure [6], as AlphaFold [7] or TCRmodel2 [8], and trying to embed it into systems predicting molecular bindings [9], even if much work is still to be done.

Many approaches proposed in the literature present good performance when input data are not strictly divided, but the same performance is not observed when trying to extrapolate on unseen epitopes [10].

Authors of [11] try to embed physical and biochemical properties in their neural network model for binding prediction, arguing that amino acid physical proximity could not have the greatest importance in determining TCR-epitope specificity, in open contrast to other works [5], [12].

The problem of improving how a specific model generalizes to unseen data is addressed in [13], which proposes a method to split data in such a way that testing is only performed on sequences not used during training, and exploits a negative sample strategy. Negative data are included in [14] as well, which also discusses the employment of pretraining and concludes that, if not well-balanced, it leads to a drop in performance. On the other hand, authors of [15] and [16] concluded that self-supervised pretraining highly improved their model performance for some specific cancer types.

A predictor for TCR-peptide binding only focusing on epitopes with enough known TCRs is presented in [17]; this work concludes that a good performance for generalization to unseen data can be obtained only for epitopes having a high similarity and the same MHC restriction to epitopes in the training set.

Also, most models in the literature are made to pair CDR3s only with known epitope peptides. However, our knowledge of epitope peptides and our databases are incomplete, which means that a classification system that can deal only with known epitopes is fundamentally flawed from its conception. The real problem is not to just pair an unseen CDR3 with an existing epitope peptide but to find what type of sequence this peptide should have in order to pair with that specific CDR3.

Besides deep learning approaches, more traditional machine learning methods have been exploited to try to solve the problem of TCR-epitope binding prediction. Physico-chemical features and negative samples are embedded, as well, in a Support Vector Machine model presented in [18], which highlights the need for considering the possible affinity among different peptides.

The great variety and diversity of the cited work in the literature and the often opposite ideas and conclusions drawn in all of them underline how the binding prediction problem is still quite far from being solved.

## 3. Methodology

### 3.1. Dataset Description

The dataset used in this study is pivotal for understanding the interactions between epitope peptides, CDR3 sequences and MHC molecules. This dataset is essential for immunotherapy techniques, particularly in the context of predicting which epitope peptide can bind to a specific CDR3 sequence when presented by a defined MHC molecule. By identifying the exact target of a specific TCR, such prediction enables the use of the given TCR in adoptive T-cell therapy approaches.

Some of the dataset entries are reported in Figure 1 and include the following columns:

- **Epitope peptide**: This column lists the amino acid sequences that define each epitope peptide. Epitope peptides are the part of antigens that are recognized by T lymphocytes, a major arm of the immune system. As highlighted by [1], these peptides are recognized through the TCR molecule and are critical in initiating T cell-mediated immune responses, with MHC class I molecules typically binding peptides of 8-10 amino acids, and MHC class II molecules binding longer peptides of 13-18 amino acids. Understanding these sequences is crucial for designing vaccines and for cancer immunotherapy, as they can trigger an effective immune response.
- **CDR3 beta aa**: This column contains the amino acid sequences of the CDR3 region of the T-cell receptor (TCR) beta chain. The TCR is composed of an alpha and a beta chain. In each chain, the CDR3 (Complementarity-Determining Region 3) is a critical part as it directly interacts with the epitope presented by the MHC molecule. As discussed by [1], the CDR3 region’s high variability is essential for antigen specificity, as it directly contacts the central portion of the peptide within the MHC groove, allowing the immune system to recognize a vast array of antigens.
- **MHC**: This column specifies the type of MHC molecule. Major Histocompatibility Complex (MHC) molecules play a crucial role in the immune system by presenting peptide fragments (epitopes) to T cells. According to [1], class I MHC molecules present antigens derived from intracellular pathogens to CD8+ cytotoxic T cells, while class II MHC molecules present antigens from extracellular pathogens to CD4+ helper T cells. This differentiation is key to facilitating a broad range of immune responses.

**Figure 1:**
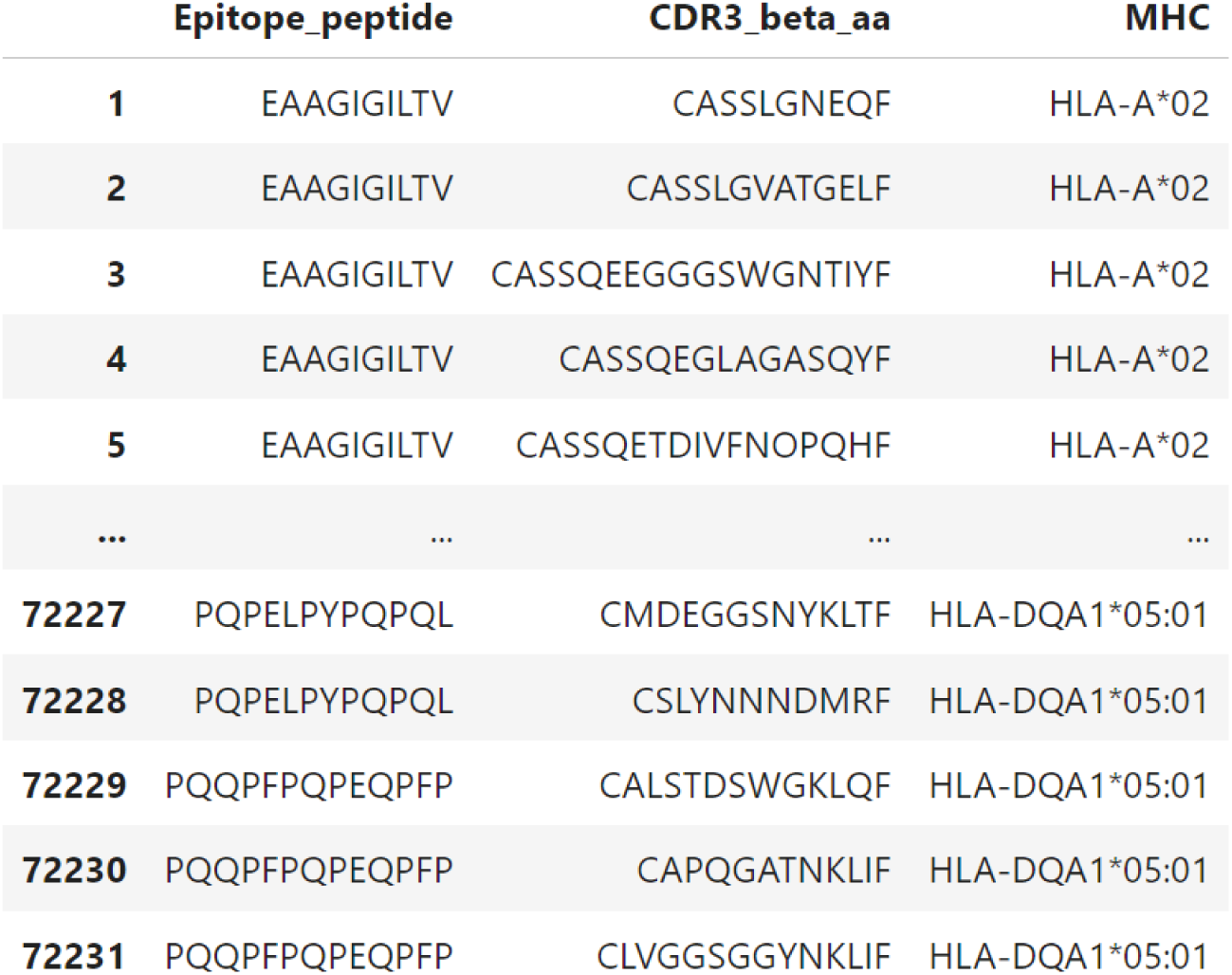
Initial and final part of the dataset

In the context of immunotherapy, and cancer immunotherapy, understanding these interactions can help in predicting which peptides are likely to be recognized by T cells, thereby enabling the design of effective therapeutic strategies. For instance, as [1] indicates, identifying epitopes that can strongly bind to both MHC and TCR can lead to the development of personalized cancer vaccines or adoptive T-cell therapy that enhances the immune system’s ability to target and destroy cancer cells.

### 3.2. Data Preprocessing

In this subsection, we describe the preprocessing steps taken to prepare the dataset for the model training phase. These steps ensured data quality and consistency, enhancing the model’s performance and reliability.

First, the datasets containing information about epitope peptides, CDR3 sequences and MHC molecules were loaded and combined. Duplicate entries and rows with missing values were removed to maintain data integrity. Additionally, any row with special amino acids in key columns was excluded to prevent inconsistencies. Ensuring data integrity at this stage is critical, as emphasized by various studies such as [19], which link preprocessing quality directly to the reliability of machine learning outcomes.

To address the imbalance in the dataset, epitopes that appeared with high frequency were downsampled to a threshold of 1% of the total entries. Conversely, rare epitopes that appeared less than 0.1% of the time were removed. This balancing act is vital to avoid model bias toward more frequent epitopes, a common issue in bioinformatics. For example, in [19], immuneML applied a similar balancing strategy when working with immune receptor repertoires to prevent the proposed models from being overly influenced by highly frequent sequences, thereby enhancing the models’ ability to generalize across a broader spectrum of data. Additionally, [20] explored balancing techniques such as 2-gram encoding, which we also tried to incorporate in some of our experiments but without significant results, and the 6-letter exchange group method to maintain a balance between computational efficiency and classification accuracy in protein sequence data.

For further preparation, all text data was standardized to uppercase, and empty strings were replaced with missing values. This standardization ensured consistency across the dataset, which is particularly important when working with sequence data. While some approaches, such as the one described in [21], focus on transforming protein sequences into detailed numerical representations, we opted for a more straightforward standardization and tokenization method, given our model’s specific requirements.

Next, the data was prepared for model training by separating the features (CDR3 sequences and MHC types) and the target variable (epitope peptides). The sequences were tokenized to convert them into numerical form and then padded to ensure uniform length across all data points. This method ensures that each sequence is properly formatted for input into the model. Although reducing computational load during the prediction phase is critical, as discussed in [22], our primary focus during preprocessing was to ensure data quality and consistency, rather than optimizing for computational efficiency at this stage.

The CDR3 sequences and epitope peptides were tokenized using character-level tokenizers. These tokenizers were fitted on the training sequences and then used to transform the validation and testing sequences. Padding was then applied to standardize the sequence lengths, a common requirement for machine learning models dealing with sequence data. By following this approach, we ensured that the model would not encounter issues related to varying input lengths, which could complicate training.

To ensure that the model could generalize to new data, the dataset was split into training, validation, and testing sets using K-Fold cross-validation. This technique, which involves creating multiple folds with distinct sets of CDR3 sequences, is a well-established method for preventing overfitting and ensuring that the model’s performance is robust. By dividing the data in this way, we could accurately assess the model’s ability to generalize to unseen data [22].

Additionally, one-hot encoding was applied to the MHC features to convert them into a numerical format suitable for model input. This encoding preserved the categorical nature of MHC types while ensuring compatibility with the machine-learning algorithms used in our model, an approach that remains a standard in the field [23], [19].

By implementing these preprocessing steps, we ensured a high-quality, balanced, and consistent dataset, ready for training a robust sequence-to-sequence model aimed at predicting epitope peptides based on given CDR3 sequences and MHC molecules.

### 3.3. Model Architecture

The core of our model is based on a deep learning approach using recurrent neural networks (RNNs), specifically Long Short-Term Memory (LSTM) layers. LSTMs are particularly well-suited for sequence data as they can effectively capture long-range dependencies and patterns within the sequences, a capability that is crucial when dealing with the sequential nature of biological data, as highlighted in the study on pMTnet [24]. This model similarly employed LSTM networks for encoding MHC-peptide sequences, demonstrating the effectiveness of such an approach in capturing the dependencies necessary for predicting TCR-pMHC interactions.

Our model architecture includes the following key components:

- **Embedding Layer**: this layer transforms the input sequences into dense vector representations, allowing the model to work with numerical data while preserving the contextual relationships between sequence elements. This approach aligns with methods discussed in [25], where embedding layers were effectively used to capture the nuances in TCR sequences and improve the model’s ability to distinguish between different antigen specificities.
- **Convolutional Layers**: two convolutional layers were added to capture local patterns and features within the sequences. These layers help in extracting relevant features before passing the data to the LSTM layers. The use of CNNs to capture local patterns in sequence data is a technique that has proven effective in other models like NetTCR-2.0 [26], where convolutional layers were pivotal in analyzing paired TCR*α* and *β* sequences, enhancing the model’s ability to predict peptide binding.
- **Bidirectional LSTM Layers**: two bidirectional LSTM layers were used to capture dependencies in both forward and backward directions. This bidirectional approach ensures that the model can learn from the entire context of the sequences, a strategy also employed in deep learning frameworks like DeepTCR [25], which utilizes such layers to analyze TCR sequence data comprehensively, improving classification performance.
- **Attention Layer**: an attention mechanism was incorporated to allow the model to focus on different parts of the input sequence when making predictions. This mechanism helps in improving the model’s ability to capture relevant features and relationships within the data. The use of attention mechanisms in models like ours is crucial, as they enable the model to weigh the importance of different sequence elements dynamically, which is particularly important in complex datasets where certain features may have more predictive power [3].
- **Dense Layers with Batch Normalization and Dropout**: several dense layers were included to further process the data, with batch normalization to stabilize and accelerate training, and dropout layers to prevent overfitting by randomly dropping units during training. These techniques are essential for maintaining the model’s performance and have been widely validated in various deep-learning architectures, including those used in immunology research [3].
- **Time-Distributed Output Layer**: the final layer is a time distributed layer that applies a dense layer to each time step of the input sequence. This layer produces the output predictions for each position in the sequence, ensuring that the model can make precise predictions across the entire length of the input.

Label smoothing was applied to enhance the generalization of the model. This technique prevents the model from becoming too confident in its predictions, thereby improving its performance on unseen data. This method is particularly useful in scenarios where the training data is limited or noisy, as it can help the model avoid overfitting to specific patterns in the data, a challenge often encountered in biological sequence prediction tasks [3].

This architecture ensures that the model can effectively learn from the provided sequences and make accurate predictions about the epitope peptides based on given CDR3 sequences and MHC molecules. The detailed structure of the model is illustrated in Figure 2.

**Figure 2:**
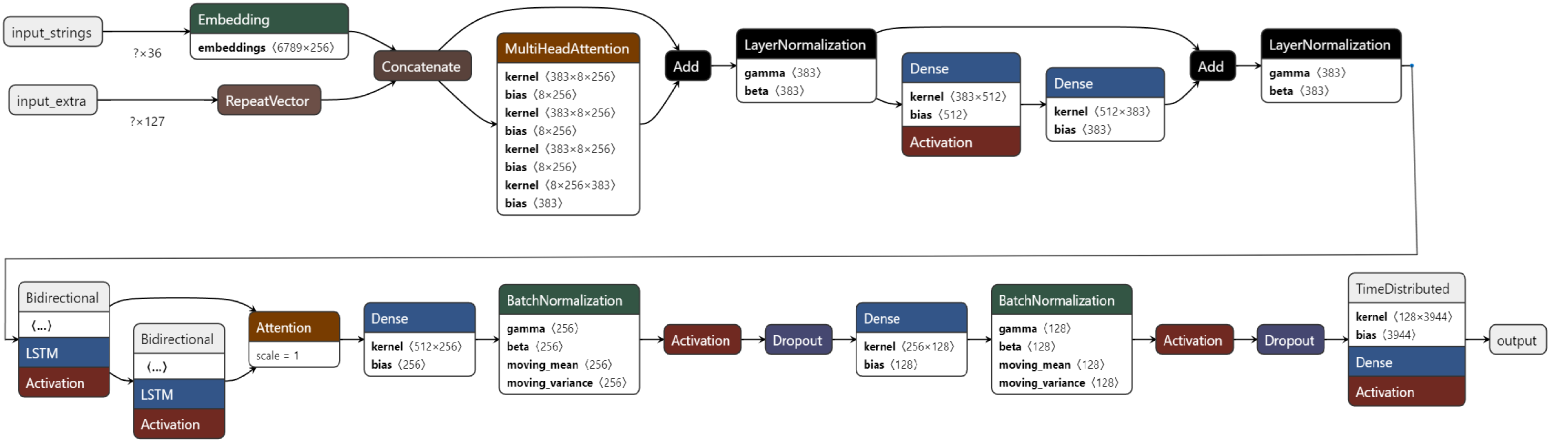
Model Architecture

### 3.4. Training Strategy

The training procedure of our sequence-to-sequence model was meticulously designed to optimize its performance in predicting epitope peptides based on given CDR3 sequences and MHC molecules. We employed several strategies to ensure effective training and evaluation of the model, drawing on best practices in deep learning and computational immunology.

The model was trained using a batch size of 64 and for a total of 200 epochs. These hyperparameters were carefully chosen based on preliminary experiments to balance between training time and model performance. The total number of training steps was calculated by multiplying the number of epochs by the number of batches per epoch, a standard approach that ensures comprehensive coverage of the dataset during training.

To further enhance the model’s learning efficiency, we implemented the OneCycleScheduler, a dynamic learning rate adjustment technique. This scheduler gradually increases the learning rate during the initial phase of training and decreases it during the later phases, promoting efficient learning and convergence. The effectiveness of dynamic learning rate adjustments is well-documented, particularly in complex models dealing with biological sequences, such as those discussed in the TCRGP study [27], which also emphasizes the importance of adapting learning strategies to optimize model performance. The optimal learning rate identified for our model was 0.01.

Several callbacks were employed to ensure robust training:

- **ModelCheckpoint**: this callback saved the best model during training based on validation loss, ensuring that we retained the most effective version of the model. The use of validation-based checkpoints is a common practice in deep learning, as it prevents overfitting and ensures that the final model is the best representation of the learning process.
- **EarlyStopping**: this callback monitored the validation loss and stopped training if no improvement was observed for 10 consecutive epochs, thereby preventing overfitting and saving computational resources. This approach is particularly valuable in scenarios with high-dimensional data, such as TCR sequence analysis, where overfitting can significantly degrade model performance, as highlighted in TCRMatch [28].
- **OneCycleScheduler**: as mentioned, this callback adjusted the learning rate dynamically throughout the training process. This technique is increasingly recognized for its ability to accelerate convergence and improve model robustness, especially in models handling complex datasets like those involving TCR sequences [27].

The model was trained on the padded CDR3 sequences and one-hot encoded MHC features, with the epitope peptides as the target output. During training, the data was shuffled at each epoch to ensure that the model did not learn any unintended patterns from the data order, a technique that aligns with best practices in sequence-based model training to avoid introducing biases.

Finally, the best model was saved during the training process and loaded at the end for evaluation on both the validation and test sets to assess its performance. This approach of saving the optimal model based on validation performance, rather than just the final model after all epochs, helps ensure that the best possible model configuration is used for predictions, as discussed in various studies [29].

### 3.5. Comparative Analysis with Existing Models and Architectures

As far as different models go, we can see the full picture of how our model compares to the rest of the literature by looking at Table 2. In particular, our approach offers advantages and innovation with respect to the existing literature concerning the following aspects:

- **Task**: In the field of TCR-epitope interaction prediction, several models have been developed. However, most existing models focus primarily on binary classification tasks, such as determining whether a binding event is likely to occur between a TCR and an epitope. This approach, while valuable, provides a limited perspective, as it only answers if the binding is possible rather than predicting what the binding sequence might look like. In contrast, our model goes beyond binary classification by generating the amino acid sequence for the potential epitope itself. This distinction is critical: instead of merely predicting compatibility, our model constructs the precise sequence of amino acids that could successfully bind to a given CDR3 sequence. By focusing on this generative task, we aim to provide a more biologically informative prediction that aligns with the needs of personalized immunotherapy, where exact sequence matching is paramount for targeted treatment.
- **Accuracy**: Most existing models, such as DeepTCR, NetTCR-2.0, and TCRGP, excel in binary classification, reporting high AUC scores (e.g., 0.9+ for DeepTCR). However, these metrics reflect an easier task—determining whether binding occurs, not predicting which amino acids participate in the interaction. Our model achieves an 85% test accuracy for predicting the exact amino acids at each position, a more challenging and biologically informative task. Therefore, while their AUC scores are higher, our model addresses a more complex problem.
- **Explainability**: Our model also differentiates itself through attention-based interpretability, allowing researchers to understand which sequence elements drive predictions. Models like TCRGP and NetTCR-2.0, while accurate, lack such transparency. This makes our model more suitable for clinical applications, where understanding the reasons behind predictions is essential.
- **Data Handling and Generalization**: Another key advantage of our approach is how we handle data imbalance through down-sampling frequent epitopes, ensuring that our model generalizes well to unseen data. Models like TCRMatch rely on curated datasets and perform poorly when confronted with novel epitopes, whereas our model is built to generalize beyond the training data, which is crucial for real-world immunotherapy.
- **Training Efficiency**: We also prioritize efficiency. Our use of the OneCycleScheduler allows for faster convergence, making our model more practical for large-scale applications compared to models like TCRGP, which rely on computationally expensive methods like Gaussian Process optimization.

**Table 2:**
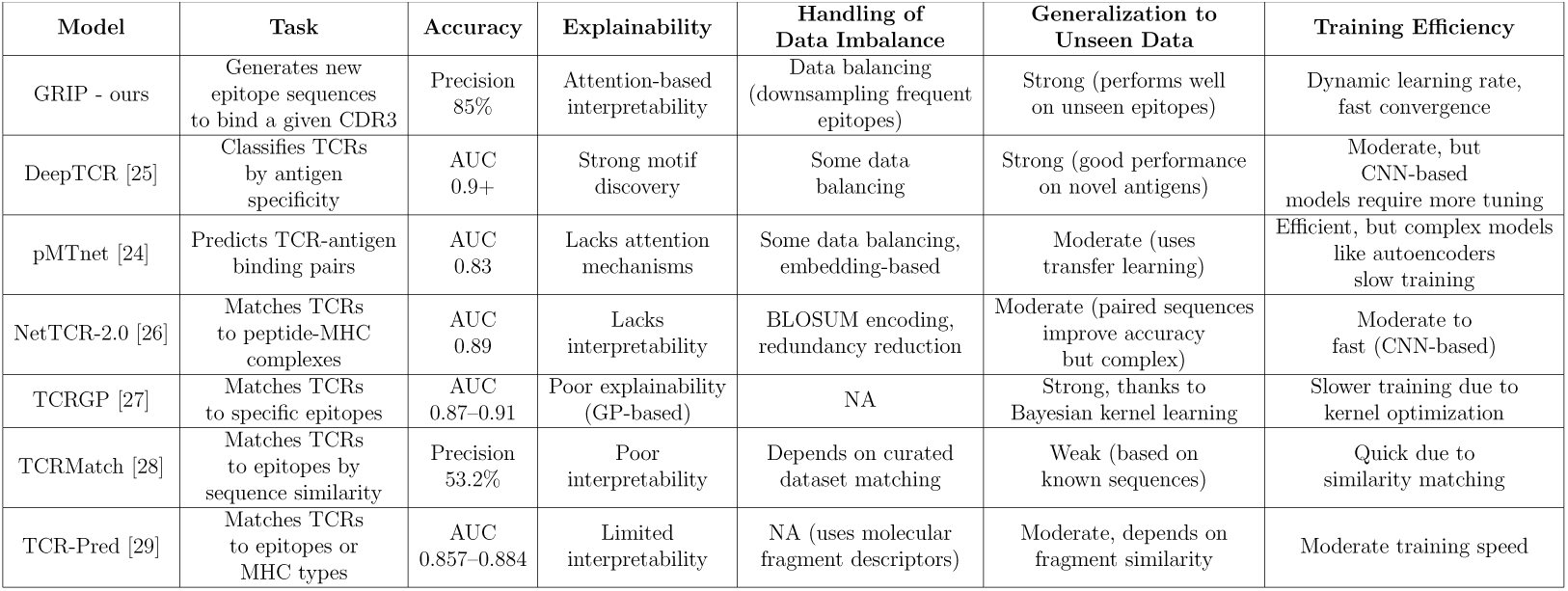
Model Comparison.

Our architectural choices are driven by the distinct demands of the generative task at hand. We chose an LSTM network over alternatives like Transformers, Convolutional Neural Networks (CNNs), or even standard RNNs and Gated Recurrent Units (GRUs) because of the LSTM’s unique suitability for handling sequential data with long-range dependencies—essential for accurately modelling the complex interactions in TCR-epitope binding.

Our model is based on a novel generative and probabilistic output approach. Unlike previous models that usually are limited to classifying between known epitopes, our model produces each amino acid in the predicted epitope sequence through a probabilistic process informed by learned patterns. At each position in the sequence, the model calculates the probability of each possible amino acid and then selects based on these probabilities. This means that each predicted epitope sequence is not a fixed, deterministic match to the training data but a new, context-driven sequence tailored to the specific CDR3 input. This generation aspect empowers our model to dynamically create sequences that are both plausible and mostly accurate, even for completely unseen CDR3 inputs.

Moreover, the attention mechanism enhances this generative process by focusing on the most relevant sequence elements, and by refining the probability distributions for each position in the generated sequence. This combination of attention-driven focus and probabilistic generation allows the model to produce contextually accurate and adaptable outputs, distinguishing it as a tool for applications where precision and flexibility are essential, such as in personalized cancer immunotherapy.

While Transformers are adept at modelling global relationships through self-attention, they can be computationally intensive, especially for long biological sequences like those in TCR-epitope interactions. CNNs, though efficient for identifying local patterns, cannot capture long-term dependencies. Additionally, standard RNNs and GRUs can struggle with vanishing gradient issues, making LSTMs an optimal choice for sustained sequential memory.

In summary, our LSTM-based model, coupled with attention mechanisms and a probabilistic generative approach, combines the benefits of sequential memory, interpretability, and dynamic sequence generation. This architecture allows us not only to predict binding but also to generate biologically meaningful sequences, positioning our model as an advanced tool for predictive and generative tasks in immunotherapy and elsewhere.

## 4. Results

### 4.1. Model Performance and Metrics

The performance of our sequence-to-sequence model was evaluated based on its ability to predict epitope peptides given specific CDR3 sequences and MHC molecules. The evaluation metrics include loss and accuracy on the train, validation and test datasets.

After training, the best model achieved the following results:

- **Train Loss**: 0.5796
- **Train Accuracy**: 97.24%
- **Validation Loss**: 0.9583
- **Validation Accuracy**: 85.83%
- **Test Loss**: 0.9768
- **Test Accuracy**: 85.16%

These metrics indicate that the model performs well in generalizing to unseen data, making it a robust tool for predicting epitope peptides.

To visualize the training process, we plotted the accuracy and loss over epochs for both the training and validation sets which helped in understanding its performance and convergence behaviour during training. More specifically, even though the validation accuracy could not reach the training accuracy, we still saw a remarkable performance for both curves and a very low loss.

To provide a more comprehensive evaluation of the model’s performance, we analyzed additional metrics beyond the basic accuracy and loss. These metrics offer a deeper understanding of the model’s strengths and areas for future improvement.

#### 4.1.1. Amino acid Level Accuracy per Position

We calculated the accuracy for each character (amino acid) at every position across the entire testing dataset. This detailed analysis helps in understanding the model’s performance for each specific amino acid at each position within the epitope peptides. The overall character-level accuracy, combining all positions, was found to be 65%.

For each position (rows), we created a detailed table, as shown in Figure 3, showing the average accuracy of every amino acid (columns) in that specific position, as well as the overall accuracy of all amino acids in that position of the peptide, see the last column (Average Accuracy). This table provides insights about which amino acids the model predicts more accurately at specific positions. Understanding these accuracies can be particularly valuable for researchers and practitioners who need precise predictions for each position in the epitope peptides. For instance, if certain positions in the peptide sequence are critical for binding to CDR3 sequences or MHC molecules, knowing the model’s accuracy at these positions can guide further research and applications.

**Figure 3:**
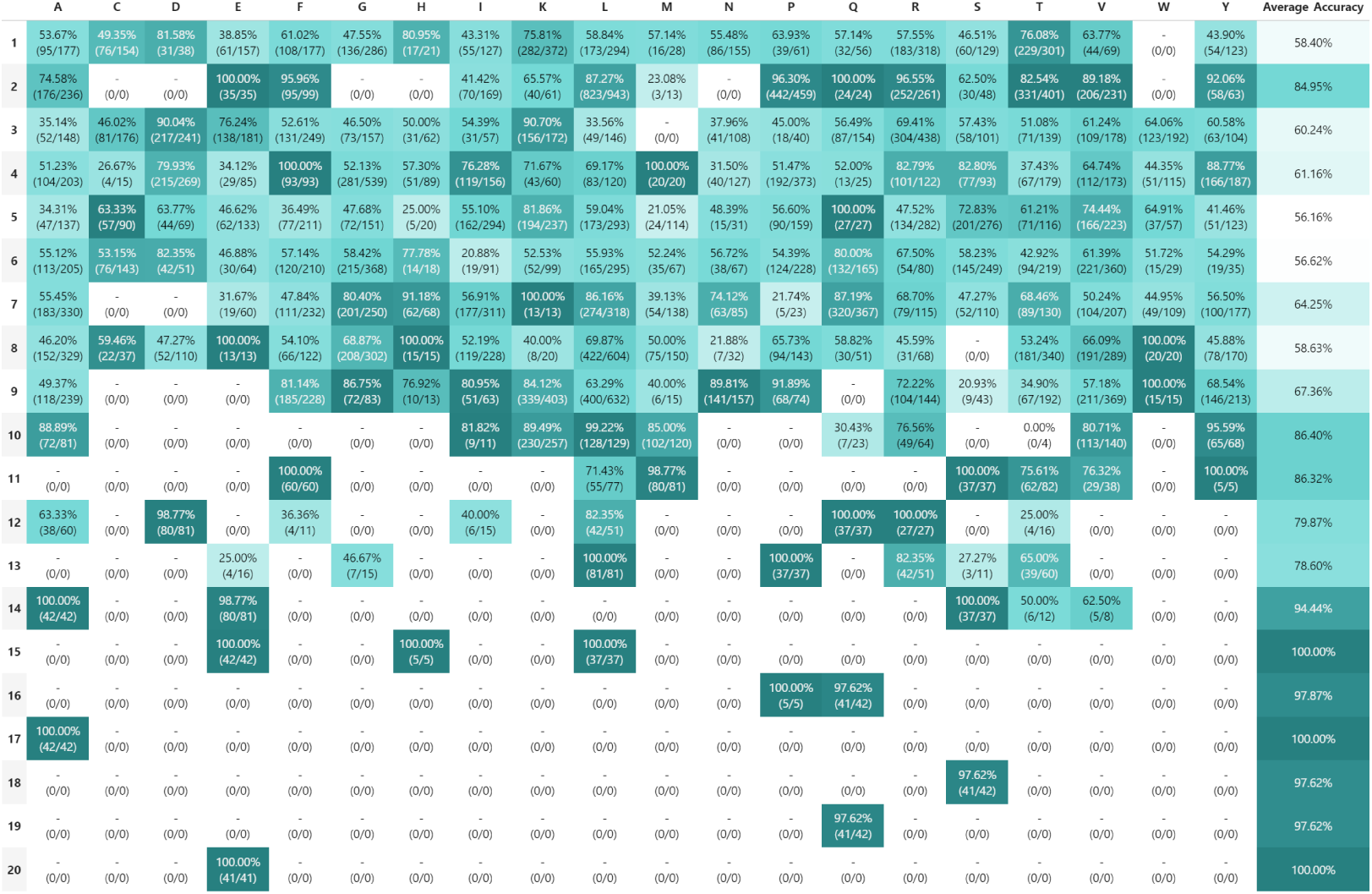
Accuracy of each amino acid in each position

#### 4.1.2. Incorrect Predictions by Position

In addition to character-level accuracy, we analyzed the incorrect predictions made by the model for each amino acid at every position. This analysis reveals which amino acids are often mispredicted and what they are typically mistaken for. This is clearly shown in Figure 4, which represents just one of all possible positions, specifically the first one. The rows represent the prediction that we make, which in this case is incorrect, and the columns are the true value, the one we should have predicted, which means that if we take A for example, in position 1 it is predicted correctly 53.67% on average, which means that 46.33% it isn’t. This 46.33% is now split into the rest of the amino acids and is presented in the cells of position 1, in row A, among all the columns. Specifically, the number one amino acid that we predict incorrectly as A is E with 10.31% and so on for every possibility.

**Figure 4:**
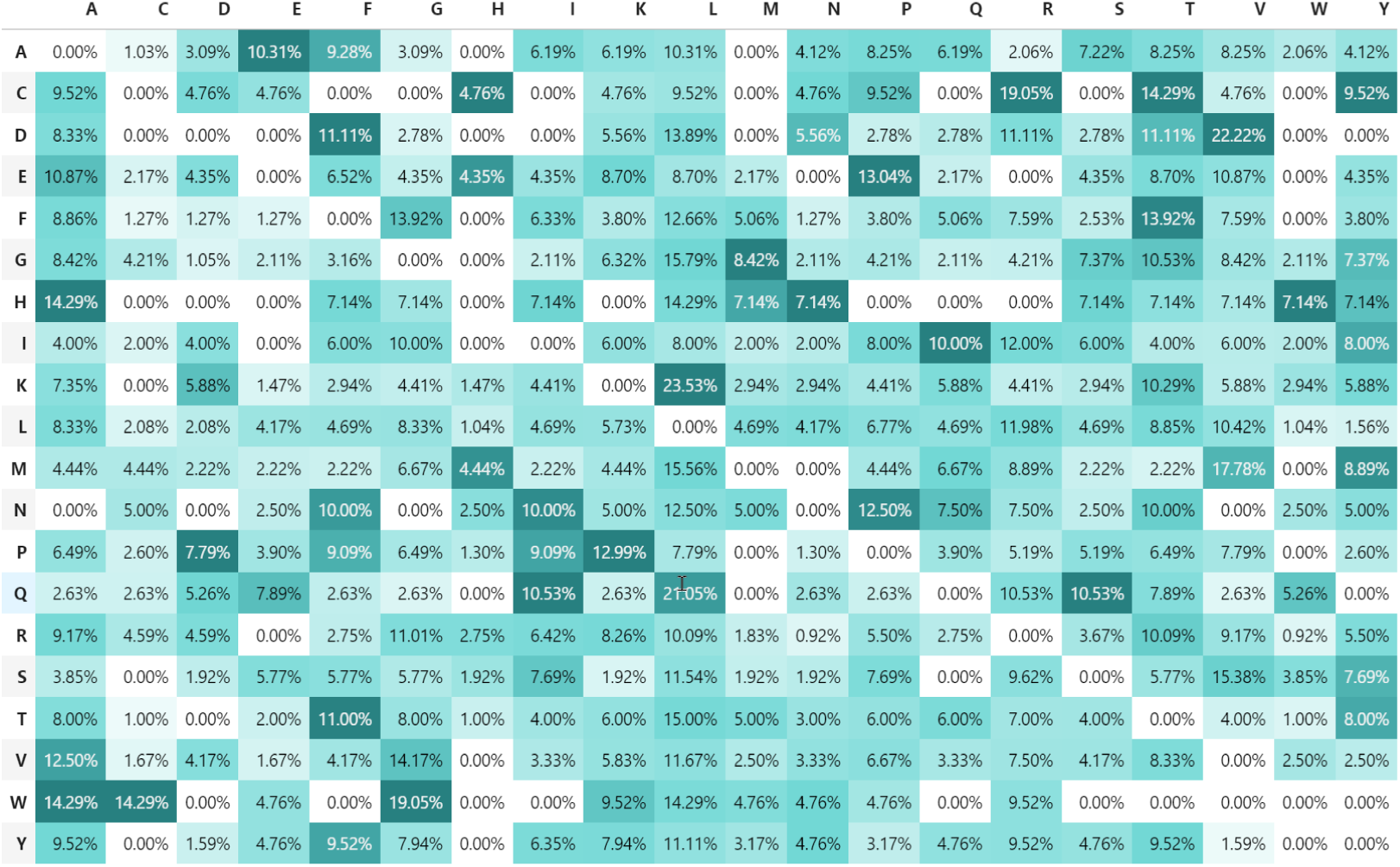
Incorrect Predictions

This information is crucial for understanding the model’s behaviour when it makes errors. By identifying common mispredictions, researchers can gain insights into potential weaknesses in the model and address these issues in future iterations. Furthermore, this analysis helps users anticipate and interpret the model’s predictions, providing a clearer picture of the model’s reliability and potential improvements.

#### 4.1.3. Epitope Frequency and Accuracy

Apart from character-wise metrics, we thought that a more zoomed-out approach would also be insightful.

As shown in Figure 5, we present the frequency and accuracy percentages of all the amino acids in the epitopes of our testing dataset. It is immediately observed that even though some epitopes appear even 10 times less than others, their overall accuracy is not always affected by their smaller frequency. On the other hand, epitopes more often appearing can potentially be predicted with a smaller accuracy; this shows that higher frequency does not correlate to higher accuracy.

**Figure 5:**
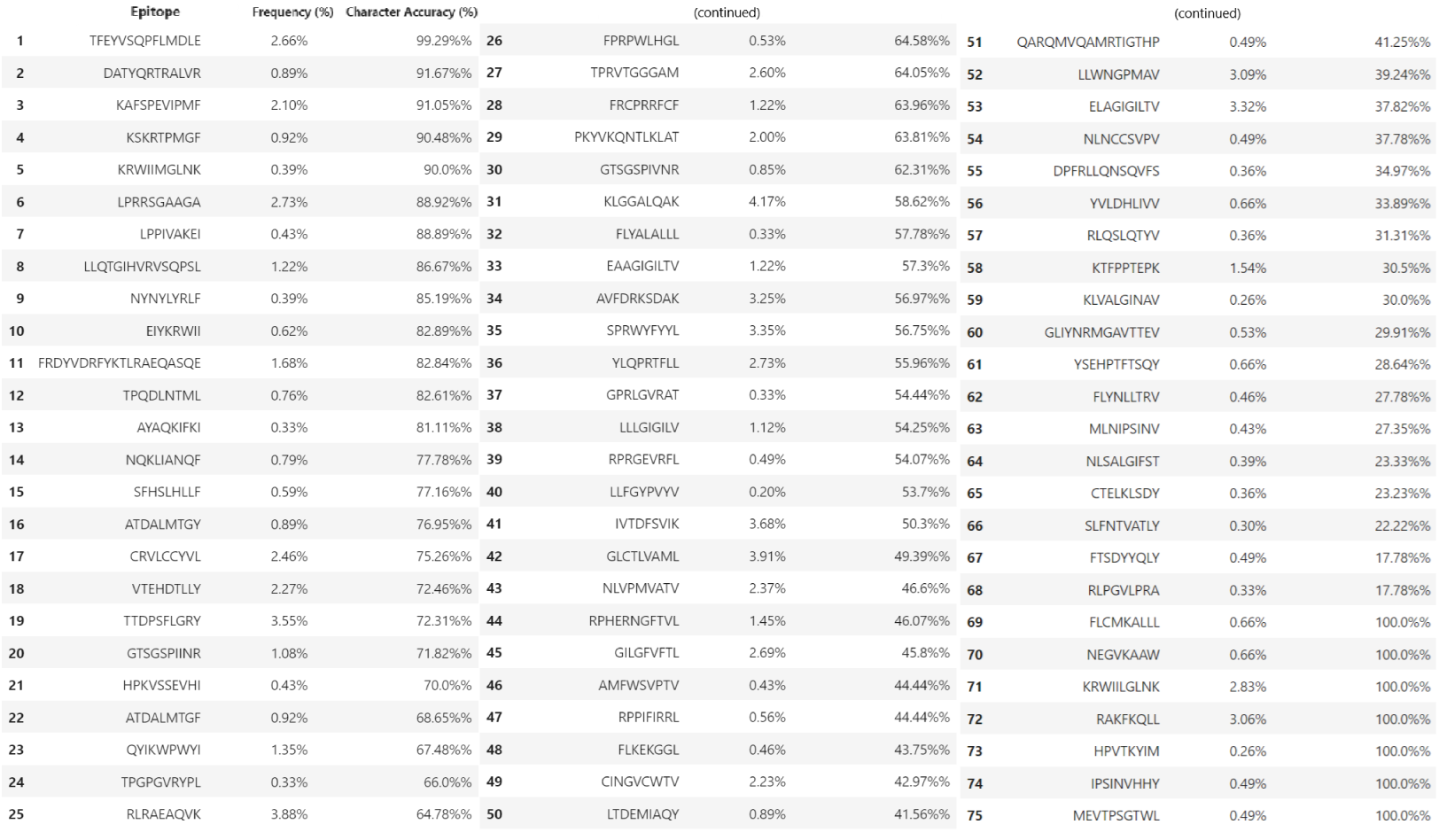
Accuracy and Frequency of all Epitopes

The combination of i) a non-over-fitting training model, ii) accuracy not bounded to frequency, and iii) different CDR3 between training, testing and validation datasets proves how unbiased and accurate our model is.

#### 4.1.4. Prediction Probabilities

The model’s binding predictions are based on the computed probabilities for each amino acid at every sequential position in the epitope peptides. To provide a more comprehensive view, we examined these probabilities for each epitope in the testing dataset. This allows users to see not only the top predictions but also the other potential predictions along with their probabilities.

This detailed probability distribution, given as an example for the first two epitopes of the testing dataset, as seen in Figure 6, can be especially beneficial for users with domain-specific knowledge. It shows exactly what probability the model gave to every amino acid in every position of all epitopes tested to be the correct prediction. By understanding the full range of probabilities, users can make more informed decisions based on the model’s evaluations, instead of just relying blindly upon the one with the highest number. For instance, in cases where the top prediction might not be the most biologically plausible, the second or third-highest probabilities might provide valuable alternatives. This level of detail empowers users to leverage the model’s predictions more effectively and apply their expertise to refine and interpret the results.

**Figure 6:**
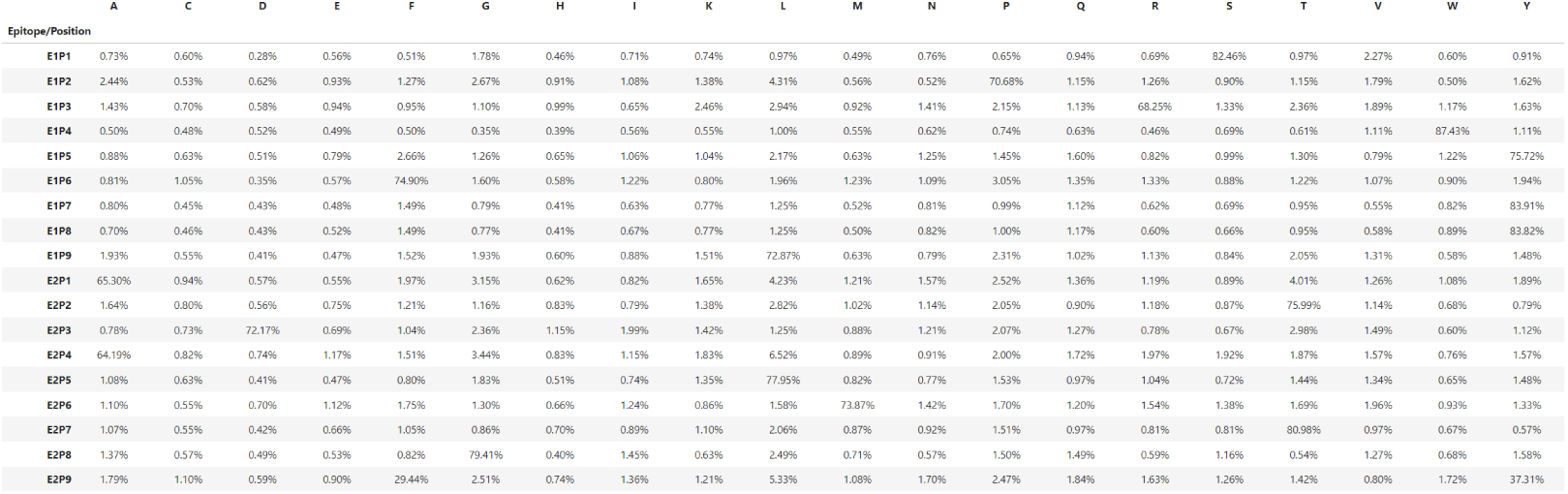
All probabilities per amino acid per position

#### 4.1.5. Multiple Predictions

Inspired by the table of probabilities for each amino acid in each position, we created a different method of predicting them. More specifically, instead of always automatically picking the amino acid with the highest probability, we give the user the freedom to select the top *N* amino acids with the best probabilities.

As we discussed before, the amino acid with the highest, given by our model, probability is occasionally not the correct prediction and there are instances where other amino acids, even with a lower estimated probability, are indeed the correct ones. Thus, we created a new way to predict accuracy which involves multiple amino acids. If the real amino acid, the one we are trying to predict is found in the group of these *N* -predicted amino acids, we count it as a correct prediction.

Using this different method, we got some interesting results where the accuracy rose significantly both overall and per position. In more detail, for each amino acid added to the total predictions, the accuracy increased based on this mathematical expression:

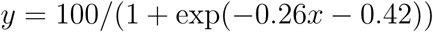

The more amino acids are included in the prediction group, the better accuracy we get in all metrics, but with a deceleration in growth. This behaviour was already predicted, but it is necessary to understand exactly when to stop including more amino acids in the prediction group. We speculate that there is a thin line between choosing a lower accuracy with fewer amino acids and higher accuracy with more amino acids, but we leave that decision to the experimentation of the user.

Overall, these detailed metrics and analyses provide a comprehensive evaluation of our sequence-to-sequence model. They highlight the model’s strengths in accurately predicting epitope peptides and provide valuable insights into its behaviour and reliability for specific amino acids and positions within the peptides. These results underscore the model’s potential for advancing personalized immunotherapy by accurately identifying epitope targets based on immune receptor sequences.

## 5. Discussion

Our research demonstrates the feasibility and effectiveness of using deep learning techniques to model complex biological interactions. The sequence-to-sequence model we developed captures the intricate dependencies between epitope peptides, CDR3 sequences, and MHC molecules. These results highlight the model’s potential as a robust tool for predicting peptide binding, thereby contributing to the advancement of targeted immunotherapy strategies.

One of the key strengths of our approach is the use of generative deep learning to address the inherent complexity of TCR-epitope binding, which allows for novel epitope predictions rather than relying solely on existing databases. Our model’s use of attention mechanisms offers a significant advantage in interpretability, enabling insights into which features most influence binding predictions. Additionally, GRIP’s ability to synthesize new epitopes, even with unseen inputs, represents an important step forward in personalized immunotherapy, potentially improving patient outcomes by providing more tailored treatment options. These innovations position our model as a valuable tool for exploring immune interactions in a way that is both predictive and generative, setting it apart from conventional methods.

Despite the promising results, there are some limitations and areas for improvement in our study.

Accuracy Improvements: While the model achieves high accuracy, there is room for improvement, particularly in reducing the test loss. This is a common challenge in deep learning models for biological data, as seen in studies like NetTCR-2.0 [26], where fine-tuning for optimal accuracy across diverse datasets remained a significant hurdle.

Data Imbalance: Although we addressed data imbalance through down-sampling and rare epitope removal, the problem persists. As noted in immuneML [19], data imbalance requires advanced solutions beyond simple downsampling. Future work could involve synthetic data generation techniques, such as SMOTE, to better balance datasets, as suggested in related studies [20].

Lack of Structural Information: The reliance on sequence data limits our model’s ability to leverage the rich structural and chemical properties of amino acids and peptides. This limitation, observed in other models like pMTnet [24], underscores the need for integrating biophysical and 3D structural data, which could enhance predictive power and biological relevance [3, 23].

Generalization Challenges: Our model shows robustness but faces difficulties in generalizing to new data, particularly unseen TCR or MHC variants. Similar challenges are noted in TCRGP [27]. Strategies like transfer learning or biology-informed neural networks could improve adaptability.

By addressing these limitations and incorporating additional layers of biological context, we aim to refine the model further, enhancing its utility for diverse immunotherapy applications.

## 6. Conclusion

This study establishes a novel framework for computationally predicting TCR-epitope binding interactions, demonstrating the power of deep learning to solve complex biological problems. Our sequence-to-sequence generative model, GRIP, goes beyond existing classification-focused approaches by generating novel epitope sequences tailored to specific CDR3 inputs, a significant advancement in the field.

The implications of this work are broad, particularly for personalized immunotherapy. GRIP’s ability to handle unseen data and generate biologically plausible epitope sequences highlights its potential for advancing adoptive T-cell therapy in cancer treatment. This capability could enable the design of more targeted and effective therapies, addressing the challenges posed by the vast diversity of potential epitopes and the limited availability of experimental data.

While our results are promising, they also underscore the challenges inherent in modeling TCR-epitope interactions, such as data imbalance, reliance on sequence-based features, and generalization to unseen variants. Overcoming these challenges will require incorporating structural and chemical information, leveraging advanced data augmentation techniques, and exploring physics-informed neural networks tailored for biological systems.

In summary, this work not only provides a strong foundation for computational immunology but also points toward exciting future directions. By addressing the outlined limitations and refining the model, GRIP could become an indispensable tool in personalized immunotherapy, paving the way for improved patient outcomes and novel insights into the immune system’s complex dynamics.

## CRediT Authorship Contribution Statement

**Athanasios Papanikolaou**: Data curation, Investigation, Methodology, Software, Visualization, Writing – original draft, Writing – review and editing

**Vladimir Sivtsov**: Data curation, Investigation, Methodology, Software, Visualization, Writing – original draft, Writing – review and editing

**Enrica Zereik**: Data curation, Investigation, Methodology, Software, Validation, Writing – original draft, Writing – review and editing

**Elliana Ruggiero**: Data curation, Investigation, Methodology, Validation, Writing – original draft, Writing – review and editing

**Chiara Bonini**: Conceptualization, Data curation, Investigation, Methodology, Supervision, Writing – original draft, Writing – review and editing

**Fabio Bonsignorio**: Conceptualization, Data curation, Investigation, Methodology, Software, Supervision, Writing – original draft, Writing – review and editing

## Ethics Statement

This research accurately reflects the work conducted, ensuring that all presented data are correct and that the methodologies are described in sufficient detail to enable replication by others.

This manuscript is entirely original, and any use of others’ work or words has been properly cited or quoted, with necessary permissions obtained where applicable.

This material has not been published, in whole or in part, elsewhere and is not currently under consideration for publication in another journal.

Generative AI and AI-assisted technologies have not been used to create or modify images.

All authors have been actively involved in the substantive work leading to this manuscript and accept joint and individual responsibility for its content.

## Acknowledgements

VS, AP and FB are funded by the European Union’s Horizon 2020 Research and Innovation Programme under the Grant Agreement No. 952275. CB is supported by AIRC (Ig 24965) and AIRC 5xMille, Rif. 22737, Italian Ministry of Research and University (PRIN 2017WC8499, 2022SLL3YZ), Italian Ministry of Health (Research project on CAR-T cells for haematological malignancies and solid tumors and RF-2019–12370243) EU (T2Evolve, Join4ATMP) RF-2021-12373598 “Uncover and overcome senescence and dysfunction of genetically engineered T lymphocytes for cancer immunotherapy”. ER is supported by 2023 DKMS John Hansen Research Grant.

